# BaRTv1.0: an improved barley reference transcript dataset to determine accurate changes in the barley transcriptome using RNA-seq

**DOI:** 10.1101/638106

**Authors:** Paulo Rapazote-Flores, Micha Bayer, Linda Milne, Claus-Dieter Mayer, John Fuller, Wenbin Guo, Pete E Hedley, Jenny Morris, Claire Halpin, Jason Kam, Sarah M. McKim, Monika Zwirek, M. Cristina Casao, Abdellah Barakate, Miriam Schreiber, Gordon Stephen, Runxuan Zhang, John WS Brown, Robbie Waugh, Craig G Simpson

**Affiliations:** Information and Computational Sciences, James Hutton Institute, Invergowrie, Dundee DD2 5DA, UK.; Biomathematics and Statistics Scotland, Aberdeen, AB25 2ZD, UK.; Cell and Molecular Sciences, The James Hutton Institute, Invergowrie, Dundee DD2 5DA, UK.; Division of Plant Sciences, School of Life Sciences, University of Dundee at the James Hutton Institute, Dundee DD2 5DA, UK.; Institute of Biological, Environmental and Rural Sciences, Aberystwyth University, Gogerddan, Aberystwyth, Ceredigion SY23 3EB, UK.; MRC Protein Phosphorylation and Ubiquitylation Unit, Sir James Black Centre, School of Life Sciences, University of Dundee, Dundee, DD1 5EH, UK.

**Keywords:** Barley, Reference Transcript Dataset, Transcriptome, Differential gene expression, Differential alternative splicing

## Abstract

**Background:** Time consuming computational assembly and quantification of gene expression and splicing analysis from RNA-seq data vary considerably. Recent fast non-alignment tools such as Kallisto and Salmon overcome these problems, but these tools require a high quality, comprehensive reference transcripts dataset (RTD), which are rarely available in plants.

**Results:** A high-quality, non-redundant barley gene RTD and database (<u>Ba</u>rley <u>R</u>eference <u>T</u>ranscripts – BaRTv1.0) has been generated. BaRTv1.0, was constructed from a range of tissues, cultivars and abiotic treatments and transcripts assembled and aligned to the barley cv. Morex reference genome (Mascher et al., 2017). Full-length cDNAs from the barley variety Haruna nijo (Matsumoto et al., 2011) determined transcript coverage, and high-resolution RT-PCR validated alternatively spliced (AS) transcripts of 86 genes in five different organs and tissue. These methods were used as benchmarks to select an optimal barley RTD. BaRTv1.0-<u>Qu</u>antification of <u>A</u>lternatively <u>S</u>pliced Isoforms (QUASI) was also made to overcome inaccurate quantification due to variation in 5’ and 3’ UTR ends of transcripts. BaRTv1.0-QUASI was used for accurate transcript quantification of RNA-seq data of five barley organs/tissues. This analysis identified 20,972 significant differentially expressed genes, 2,791 differentially alternatively spliced genes and 2,768 transcripts with differential transcript usage.

**Conclusion:** A high confidence barley reference transcript dataset consisting of 60,444 genes with 177,240 transcripts has been generated. Compared to current barley transcripts, BaRTv1.0 transcripts are generally longer, have less fragmentation and improved gene models that are well supported by splice junction reads. Precise transcript quantification using BaRTv1.0 allows routine analysis of gene expression and AS.

## Background

Barley is an important cereal crop grown across a geographical range that extends from the Arctic Circle to the hot and dry regions of North Africa, the near east and equatorial highlands. Adaptation of barley to very different growing conditions reflects important characteristics of genomic and transcriptomic diversity that leads to the success of the crop at different latitudes (Ashoub et al., 2018; Dawson et al., 2015; Russell et al., 2016). Changes in gene expression during development and in response to daily and seasonal environmental challenges and stresses drive re-programming of the barley transcriptome (Janiak et al., 2018; Ren et al., 2018; Kintlová et al., 2017; Calixto et al., 2016; IBSC, 2012). Transcriptomes consist of complex populations of transcripts produced through the co-ordinated transcription and post-transcriptional processing of precursor messenger RNAs (pre-mRNAs). Alternative splicing (AS) of pre-mRNA transcripts is the main source of different transcript isoforms that are generated through regulated differential selection of alternative splice sites on the pre-mRNA and up to 60-70% intron-containing plant genes undergo AS (Marquez et al., 2012; Mastrangelo et al., 2012; Staiger and Brown, 2013; Carvalho et al., 2013; Chamala et al., 2015; Filichkin et al., 2015; Capovilla et al., 2015; Calixto et al., 2016; Laloum et al., 2018; Szakonyi & Duque, 2018). The two main functions of AS are to increase protein diversity and regulate expression levels of specific transcripts by producing AS isoforms that are degraded by nonsense mediated decay (NMD) (Nilsen and Gravely, 2010; Kalyna et al., 2012; Reddy et al., 2013; Staiger and Brown, 2013; Lee and Rio, 2015). Extensive AS has been reported in barley (IBSC, 2012; Panahi et al., 2015; Calixto et al., 2016; Zhang et al., 2016a; Zhang et al., 2016b) and allelic diversity further contributes to the landscape of AS transcript variation among genotypes through elimination and formation of splice sites and splicing signals (Shirasu et al., 1999; Guo et al., 2014; Liu et al., 2015).

Although RNA-seq is the current method of choice to analyse gene expression, major problems exist in the computational assembly and quantification of transcript abundance from short read data with widely used programs. Such assemblies are typically inaccurate because first, they generate a large proportion of mis-assembled transcripts and second, they fail to assemble thousands of real transcripts present in the sample dataset (Pertea et al., 2015; Hayer et al., 2015). In contrast, non-alignment tools such as Kallisto and Salmon (Patro et al., 2017; Bray et al., 2016) provide rapid and accurate quantification of transcript/gene expression from RNA-seq data. However, they require high quality, comprehensive transcript references, which are rarely available in plants (Brown et al., 2017). In barley, RNA-seq data from eight different barley organs and tissues from the variety Morex, a six-rowed North American cultivar, was used to support annotation of the first barley genome sequence (ISBC, 2012). The subsequent release of the barley pseudogenome, estimated to contain 98% of the predicted barley genome content, has 42,000 high-confidence and 40,000 low-confidence genes and ca. 344,000 transcripts (Mascher et al., 2017). However, detailed analysis of individual gene models in the pseudogenome shows that the current annotation contains a high frequency of chimeric and fragmented transcripts that are likely to negatively impact downstream genome-wide analyses of differential expression and AS. In Arabidopsis, a diverse, comprehensive and accurate Reference Transcript Dataset (AtRTD2), was constructed from short read RNA-seq data by assembling transcripts with the assembly functions of Cufflinks and Stringtie, followed by multiple stringent quality control filters. These filters removed poorly assembled transcripts (e.g. with unsupported splice junctions), transcript fragments and redundant transcripts, all of which affected the accuracy of transcript quantification by Salmon/Kallisto (Zhang et al., 2015; Zhang et al., 2017a). AtRTD2 has been used for genome-wide differential expression/differential AS to identify novel regulators of the cold response and splicing factors that regulate AS in innate immunity and root development (Zhang et al., 2017b; Calixto et al., 2018; Bazin et al., 2018; Calixto et al., 2019).

Here, we describe the development of a first barley reference transcript dataset and database (<u>Ba</u>rley <u>R</u>eference <u>T</u>ranscripts – BaRTv1.0) consisting of 60,444 genes and 177,240 non-redundant transcripts. To create BaRTv1.0, we used 11 different RNA-seq experimental datasets representing 808 samples and 19.3 billion reads that were derived from a range of tissues, cultivars and treatments. We used high-resolution RT-PCR results to optimise parameters for transcript assembly and to validate differential AS in five different barley organs and tissues. We further compared the BaRTv1.0 transcripts to 22,651 Haruna nijo full-length (fl) cDNAs (Matsumoto et al., 2011) to assess the completeness and representation of the reference transcript dataset. As in Arabidopsis, we also generated a version of the RTD specifically for *<u>qu</u>*antification of *<u>a</u>*lternatively *<u>s</u>*pliced *i*soforms (BaRTv1.0-QUASI) for accurate expression and AS analysis, which overcomes inaccurate quantification due to variation in the 5’ and 3’ UTR (Zhang et al., 2017a; Soneson et al., 2019). Finally, we used BaRTv1.0-QUASI to explore RNA-seq data derived from five diverse barley organs/tissues identifying 20,972 differentially expressed genes and 2,791 differentially alternatively spliced genes amongst the samples.

## Results

### Transcript assembly and splice site determination

To maximise transcript diversity in the barley RTD assembly we selected barley Illumina short read datasets that covered different barley varieties, a range of organs and tissues at different developmental stages and plants/seedlings grown under different abiotic stresses. The datasets represent 11 different RNA-seq experiments, containing 808 samples and 19.3 billion reads (Supplementary Table S1). Most samples consisted of paired-end reads that were between 90 and 150 bp in length and produced from Illumina HiSeq 2000 or Illumina HiSeq 2500 runs. Exceptions were the dataset from Golden Promise anthers and meiocytes, which contained over 2 billion paired end 35-76 bp reads and the internode dataset which contained unpaired 100bp reads (Supplementary Table S1). The raw RNA-seq data of all samples were quality controlled, trimmed and adapters removed using FastQC and Trimmomatic (Figure 1; Supplementary Table S1). Reads were mapped to the reference genome sequence of barley cv. ‘Morex’ (Hv_IBSC_PGSB_v2) (Mascher et al., 2017) using STAR (Spliced Transcripts Alignment to a Reference) software (Dobin et al., 2013; Dobin and Gingeras, 2016) (Figure 1). To improve mapping accuracy and filter out poorly supported splice junctions from the sequence reads, while also considering the variability of expression levels in the different samples, we performed a three-pass STAR mapping. This was based on a two-pass alignment method to increase splice junction alignment rate and sensitivity by performing a high-stringency first pass with STAR, which was then used as annotation for a second STAR pass at a lower stringency alignment (Veeneman et al., 2016). We also performed a less stringent third pass with STAR to capture further splice junction read number evidence from the range of barley datasets that included different cultivars and landraces, which will show sequence variation among reads and affect their mapping. The third pass did not allow any additional splice junctions to be generated that were not already present after the second pass. The advantage of the third pass was to allow more reads to map to the splice junction and increase support for rarer splice site selections and increase transcript diversity. (See materials and Methods). The number of uniquely mapped reads after the three STAR passes ranged from 73% to 85% (data not shown) across the 11 samples. This iterative alignment and filtering process using STAR produced a robust splice junction reference dataset of 224,654 splice junctions that was used to support the identification of multiple transcripts per gene.

**Figure 1.**
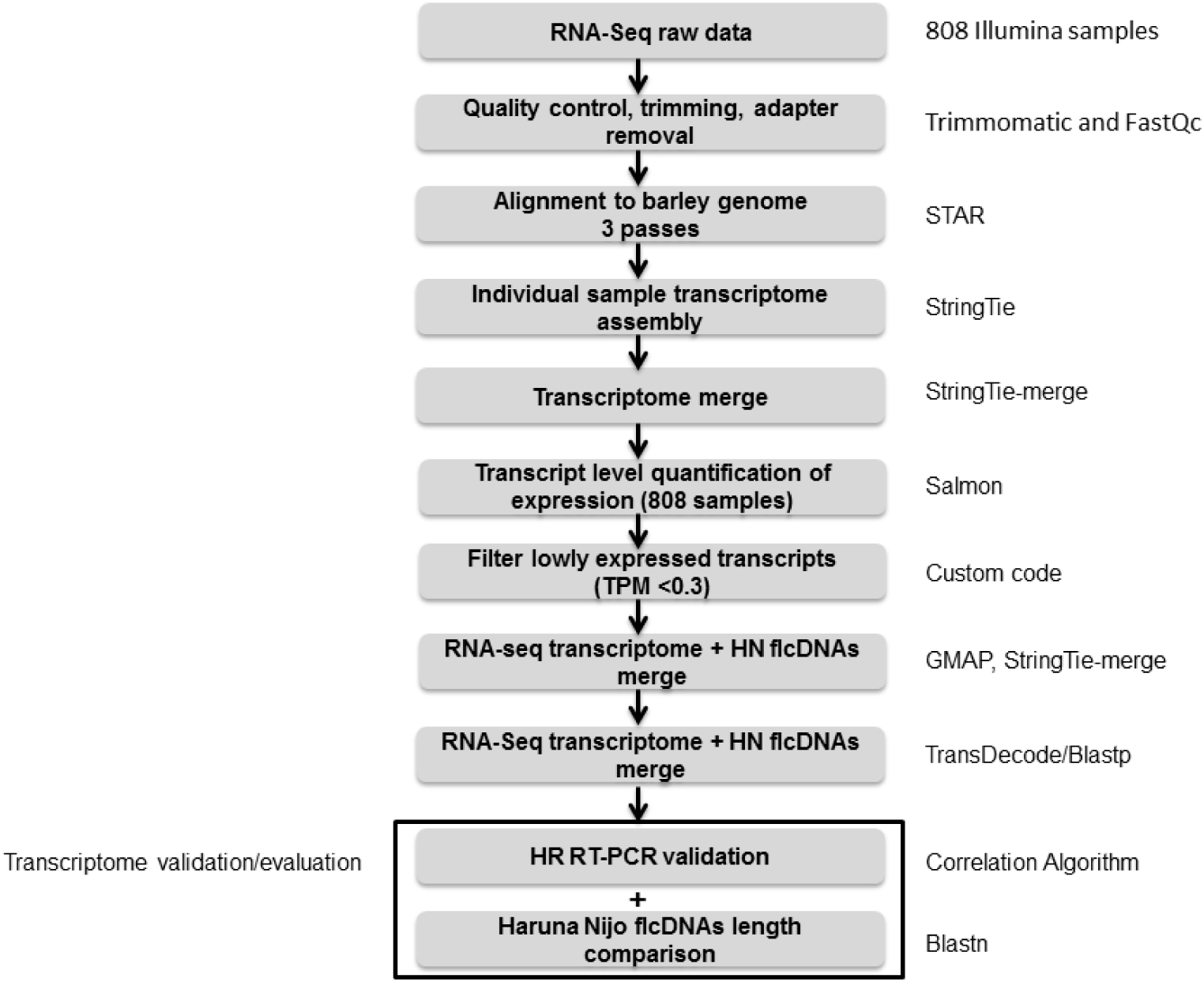
BaRTv1.0 assembly and validation pipeline. Steps in construction and validation of BaRTv1.0 and programs used in each step (right hand side).

### Optimisation of cv. Morex Guided Reference Transcript Assemblies

Transcriptomes for each of the 808 samples were assembled using StringTie (Pertea et al., 2015) and different parameter combinations tested to optimise the quality and number of transcripts (Figure 2; Supplementary Table S2). Throughout this process the quality of the Morex reference-based transcript assemblies were benchmarked against data from a high-resolution (HR) RT-PCR panel of 86 primer sets (Simpson et al., 2008) to accurately analyse the proportion of alternatively spliced products in a subset of the cv. Morex experimental samples (Developing inflorescences INF1 and INF2, leaf shoots from seedlings - LEA, embryo - EMB, internode - NOD – see Materials and Methods). The primer list is available at https://ics.hutton.ac.uk/barleyrtd/primer_list.html (Supplementary Table S3). At each stage the spliced proportions from HR RT-PCR were compared to the spliced proportions of the same AS event(s) derived from the Transcripts Per Million (TPM) counts extracted from the RNA-seq data analysis (Simpson et al., 2008; Zhang et al., 2017a) using an automated method (see Figure 1; Materials and Methods for description and https://github.com/PauloFlores/RNA-Seq-validation for script).

**Figure 2.**
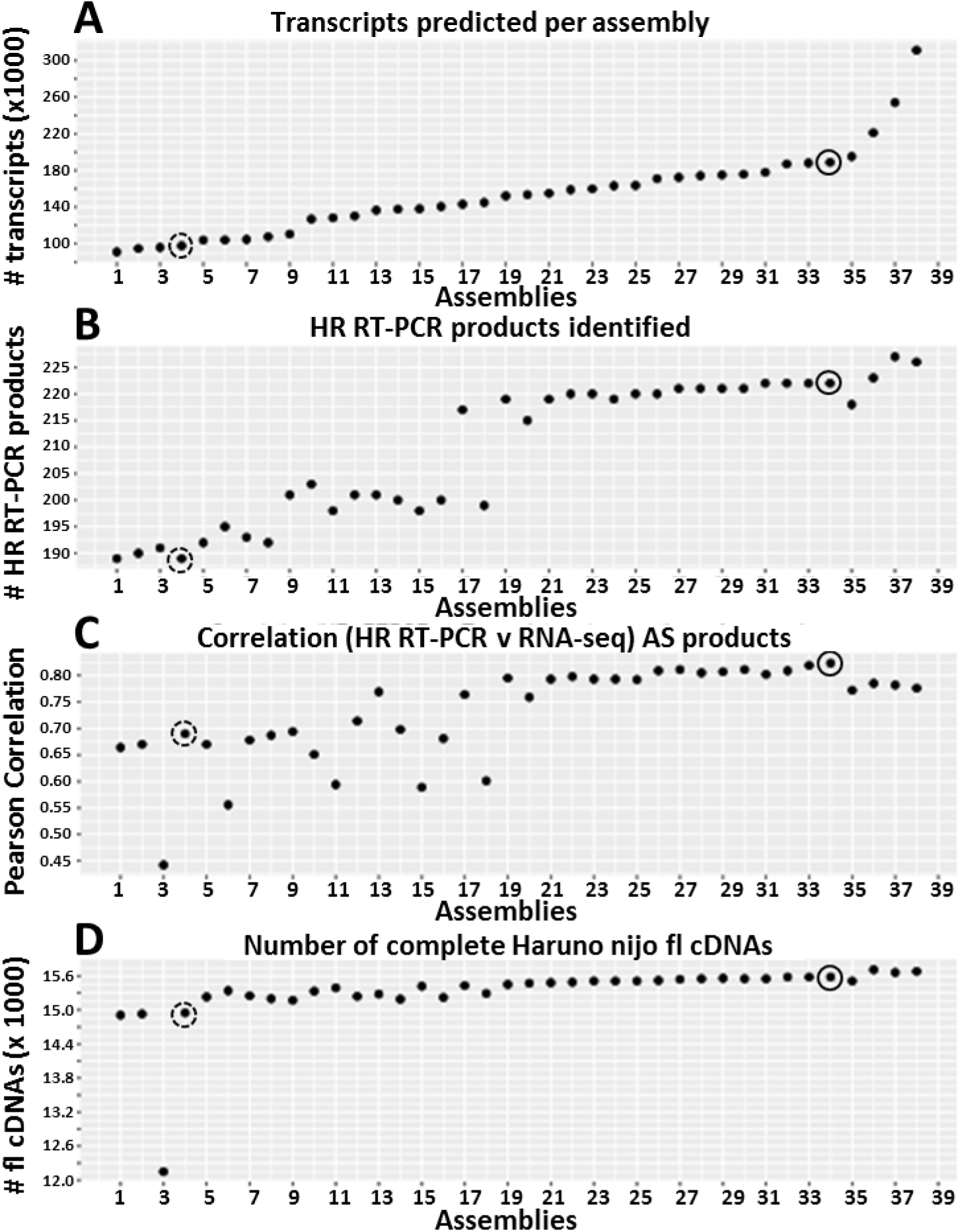
Benchmarking of 38 different StringTie Morex reference-based assemblies. The four plots show different benchmark tests to assess the parameters used in the StringTie assemblies. A) Transcript number; B) the number of HR RT-PCR products that match transcripts; C) correlation of the proportions of transcripts in 86 AS events derived from HR RT-PCR and the RNA-seq data using the different assemblies as reference for transcript quantification by Salmon; and D) the number of Haruna nijo fl cDNAs that match RTD transcripts. Each plot point represents the result of a StringTie assembly using different parameters (Supplementary Table S2). The broken circled plot points at assembly 4, an assembly using STAR defaults (without splice junction filtering) and StringTie defaults. The solid circled plot point at assembly 34 represents the selected optimised StringTie parameters used to produce BaRTv1.0 (see also Materials and Methods; Supplementary Figure 3; Supplementary Table S2).

Each StringTie assembly was further compared to the 22,651 Haruna nijo full-length (fl) cDNAs (Matsumoto et al., 2011) to assess both the completeness and representation. Of these, 17, 619 (81.2%) fl cDNAs had at least 90% coverage and 90% sequence identity with transcripts in the RTD using BLASTn (Altschul et al., 1990) (Supplementary Figure 2). These fl cDNAs were used to quantify coverage in the optimisation of assemblies with StringTie (Figure 2; Supplementary Table S2).

Among the different StringTie parameters tested, the read coverage (“-c” parameter) was found to be important and a value of 2.5 was selected as the optimum. A lower read coverage value induced fragmentation, greatly increasing the number of genes, fewer matching RT-PCR products, poorer correlation with the HR RT-PCR data and reduced matching to the Haruna nijo fl cDNAs (Figure 2, for example assemblies 9-16; Supplementary Table S2), while a value of 3 led to a lower number of genes and transcripts being defined (Figure 2, for example assemblies 26-30; Supplementary Table S2). The isoform-fraction (“-f” parameter) was optimal at 0, maximising the number of transcripts, while still maintaining a strong correlation with HR RT-PCR data and high numbers of matching Haruna nijo fl cDNAs (Figure 2, assemblies 17, 19-38; Supplementary Table S2). A minimum locus gap separation value (“-g” parameter) of 50 bp was selected as an optimum value. Values greater than 50 bp led to the prediction of fewer transcripts and poorer correlation with the HR RT-PCR data, although there was a small improvement in the coverage of the Haruna nijo fl cDNAs. Increasing the gap separation to 500 bp forced distinct genes to merge resulting in longer transcripts, poorer similarity with Haruna nijo fl cDNAs and very poor correlation with the HR RT-PCR data due to the creation of chimeric genes (Figure 2; in assembly 3). The improvement in the assemblies with the optimised StringTie parameters is illustrated by comparison to the assembly produced using StringTie default parameters (Figure 2). The optimised assembly had an 14% increase in splice product detection in the HR RT-PCR analysis (220 versus 189 RT-PCR products) and increased Pearson correlation values from 0.60 to 0.79 between the RNA-seq data and HR RT-PCR data. It also recovered 634 more complete Haruna nijo fl cDNAs compared to the StringTie assembly run in default mode.

### Construction of BaRTv1.0

Having established optimal assembly parameters, to construct the RTD, transcripts were merged to create a single set of non-redundant transcripts. The dataset was filtered to remove poorly expressed transcripts (< 0.3 TPM) and then merged with the genome-mapped Haruna nijo full-length cDNAs (Figure 1). Finally, we used TransDecoder (Haas et al., 2013) to predict protein coding regions and BLASTp (Altschul et al., 1990) to filter out transcripts equal to or less than 300 bps long (8,767 transcripts) and showing less than 70% protein coverage and identity with the *Poaceae* reference protein dataset (Figure 1), which removed all but 25 transcripts of less than 300 bp (Supplementary Figure 4). After merging and filtering, we retained 224,654 unique splice junctions, 60,444 genes and 177,240 transcripts to establish the non-redundant reference transcript dataset named BaRTv1.0 (Table 1).

**Table 1.**
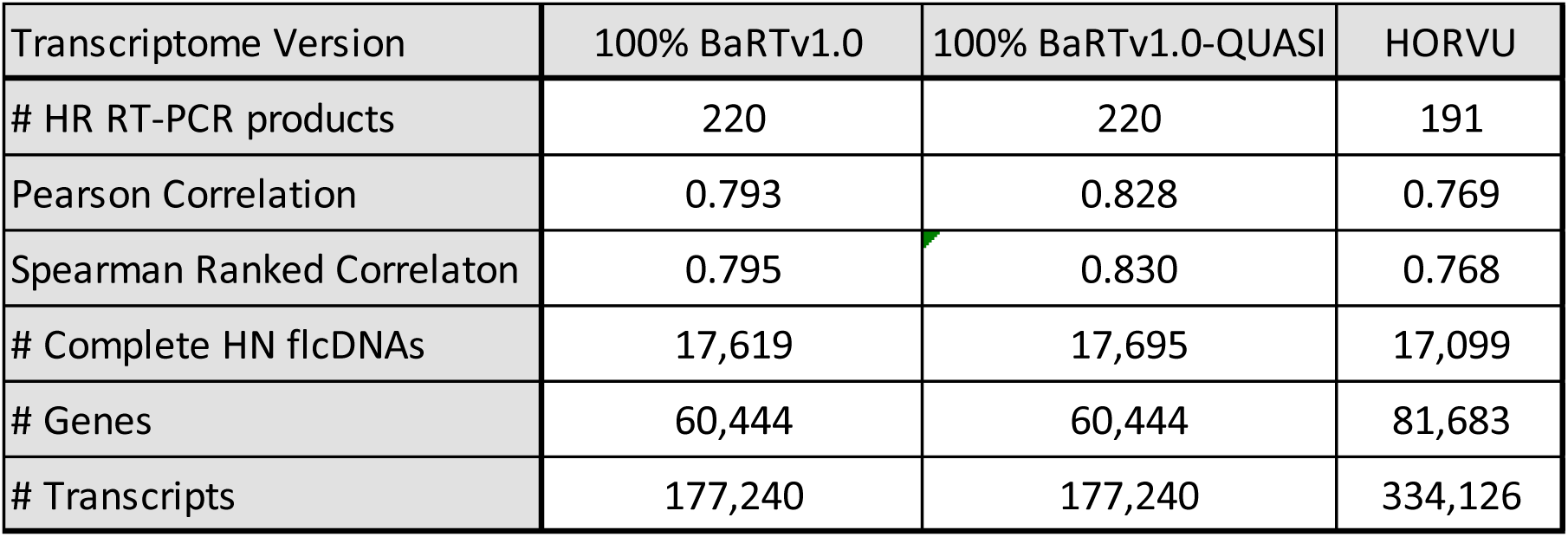
Transcriptome dataset comparisons with HR RT-PCR and Haruna nijo fl cDNAs

Previous studies in Arabidopsis and human RNA-seq analysis showed that variation in the 5’ and 3’ ends of assembled transcript isoforms of the same gene affected accuracy of transcript quantification (Zhang et al., 2017a; Soneson et al., 2019). This was overcome by padding shorter 5’ and 3’ ends to the 5’ and 3’ ends of the longest gene transcript (Zhang et al., 2017). We similarly modified BaRTv1.0 to produce transcripts of each gene with the same 5’ and 3’ ends to generate BaRTv1.0-QUASI specifically for transcript and AS quantification. Both datasets are available for download from https://ics.hutton.ac.uk/barleyrtd/downloads.html. In addition, a website was created to visualise individual BaRT transcripts, access transcript sequences, and allow for BLAST searching and comparison with existing HORVU transcripts (Mascher et al., 2017) https://ics.hutton.ac.uk/barleyrtd/index.html.

### BaRTv1.0 represents an improved barley transcript dataset

The barley cv. Morex pseudo-molecule sequences were accompanied by a set of ca. 344 k HORVU transcripts (Mascher et al., 2017), nearly double the number in BaRTv1.0. Close inspection of the HORVU transcripts identified short, fragmented and redundant transcripts. The quality control filters used in the construction of BaRTv1.0 aimed to reduce the number of transcript fragments and redundancy as these negatively impact the accuracy of transcript quantification (Zhang et al., 2017a). The BaRTv1.0 and HORVU datasets were directly compared with the numbers of complete Haruna nijo fl cDNAs and correlating the proportions of AS transcript variants measured by HR RT-PCR with those derived from the RNA-seq analysis (Supplementary Table S4). The BaRTv1.0 transcript dataset identified more of the experimentally determined HR RT-PCR products (220 versus 191) and has higher Pearson and Spearman correlations with quantification of the AS events when compared to the HORVU dataset (Table 1). For the AS events detected in BaRTv1.0 and HORVU, we plotted the percentage spliced in (PSI) values (proportion of total transcripts that include exon sequence in the transcript) from HR RT-PCR and RNA-seq for each of the three biological replicates from five different barley organ and tissue samples (giving 1992 and 1642 data points respectively) (Figure 3A and B). Pearson correlations and Spearman ranked order correlations with the HR RT-PCR data increased between HORVU (0.769 and 0.768), BaRTv1.0 (0.793 and 0.795) and BaRTv1.0-QUASI (0.828 and 0.83) (Table 1; Supplementary Table S4). We conclude that BaRTv1.0 (and the derived BaRTv1.0-QUASI) RTD is a comprehensive, non-redundant dataset suitable for differential gene expression and AS analyses.

**Figure 3.**
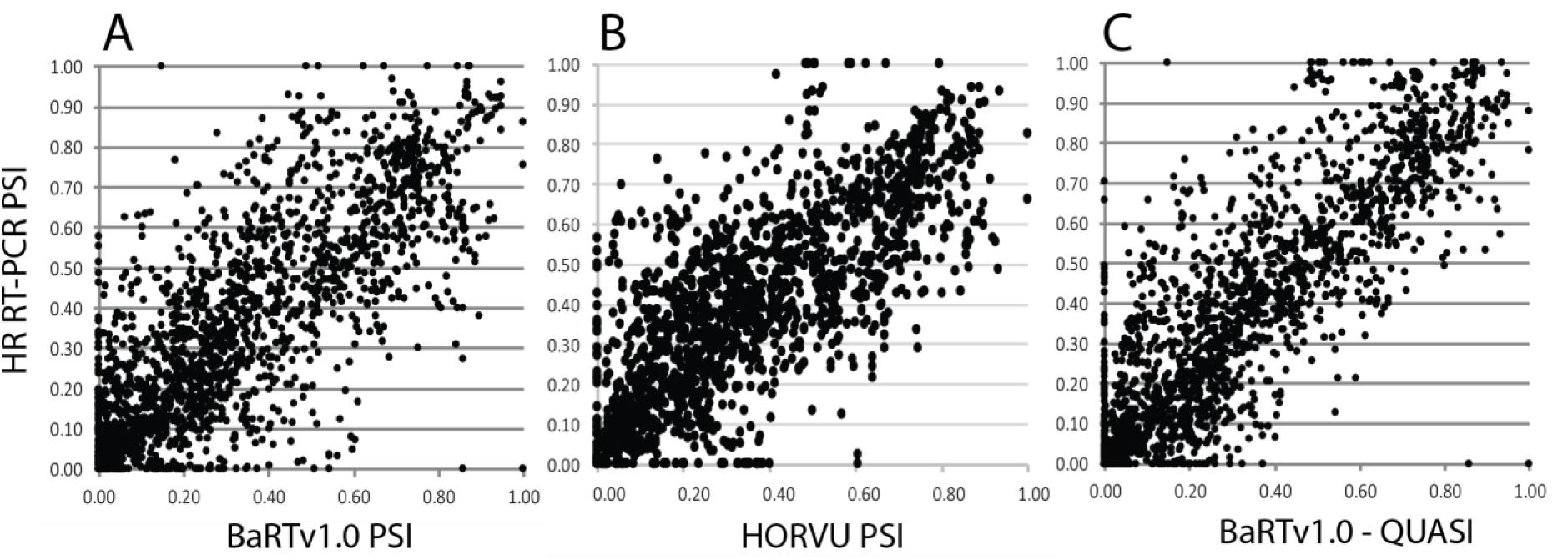
Correlation of alternative splicing from HR RT-PCR and RNA-seq. PSIs were calculated from relative fluorescence units from HR RT-PCR and transcript abundances (TPM) from RNA seq data quantified with Salmon using the A) BaRTv1.0, B) HORVU and C) BaRTv1.0-QUASI transcript datasets as reference. The 86 primer pairs designed to cv. Morex genes covered 220 AS events in BaRTv1.0 (three biological replicates of 5 different barley organs/tissues) giving 1,992 data points and 81 primer pairs covered 191 AS events giving 1,642 points for HORVU.

### BaRTv1.0 genes and transcripts

We next explored the characteristics of BaRTv1.0 genes and transcripts. A total of 57% of the BaRTv1.0 genes contained introns and had on average ∼7.7 exons per gene (Table 2). Around 60% of the multi-exon genes had multiple transcripts supporting the occurrence of widespread AS in barley. Each transcript isoform within the dataset is unique based on splice site usage (containing at least one unique splice site). Analysis of the 177,240 predicted transcripts in BaRTv1.0 showed the expected distribution of canonical splice site dinucleotides. Of the 224,654 splice junctions examined, 98.2% of the introns spliced out have the expected GT..AG splice site dinucleotides, 1.7% had GC-AG dinucleotide borders, and 0.1% showed the U12-intron-dependent splicing AT-AC dinucleotide splice sites. Half of these splice junctions were observed in all the RNA-seq datasets tested but, 1.3% were unique to a single dataset, indicating unique tissue or condition specific splicing (Supplementary Table S5).

**Table 2.**
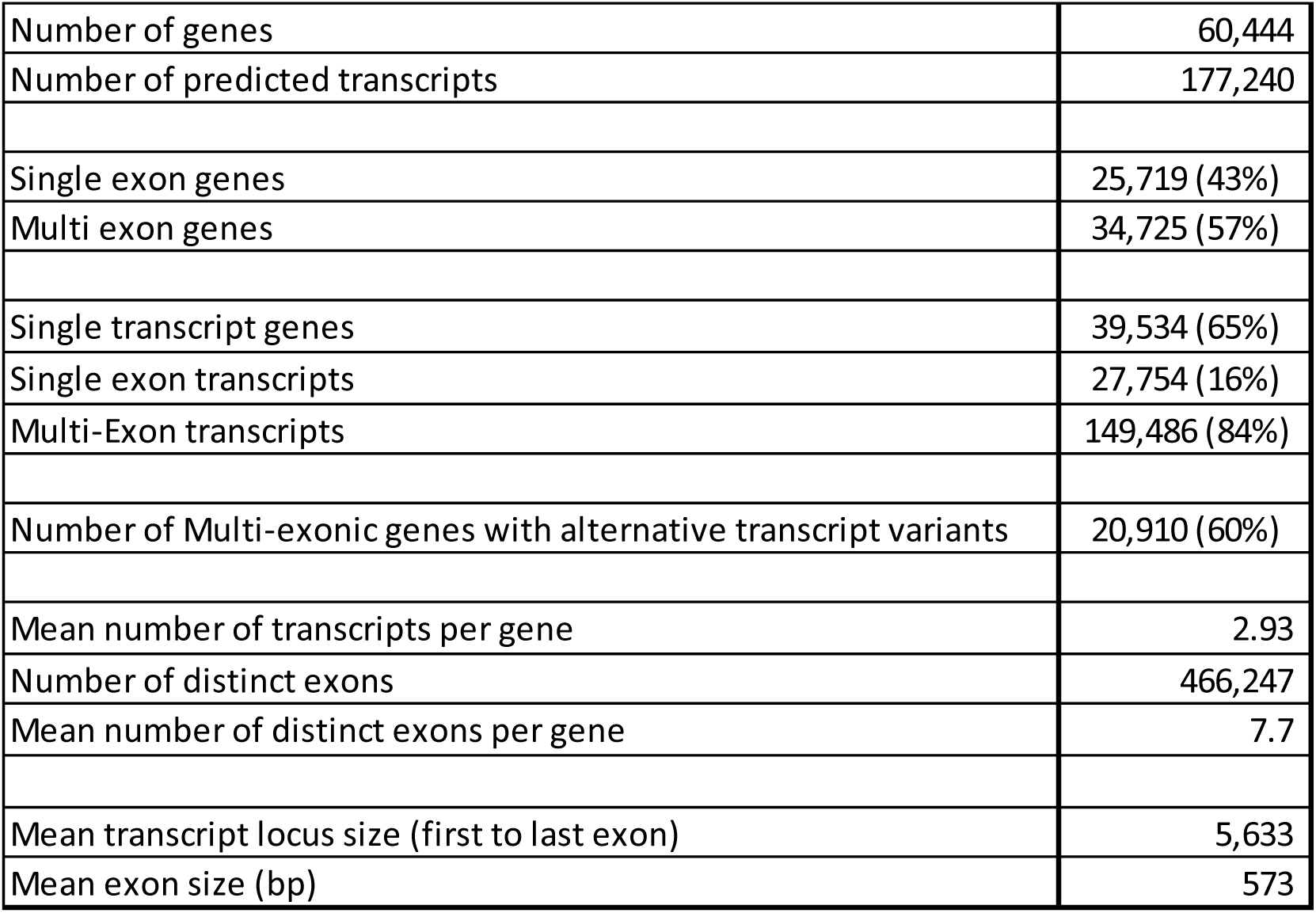
Characteristics of barley genes and transcripts in BaRTv1.0

We then used the SUPPA software version 2.3 (Alamancos *et al*, 2015) to determine different splicing events and their frequency in our transcript dataset. We identified all the expected major forms of AS, including alternative 5’ and 3’ splice site selection (Alt 5’ss; Alt 3’ss), exon skipping (ES) and intron retention (IR). Frequencies of the different AS events were consistent with studies in other plant species (Alt 5’ – 23.6%; Alt 3’ – 28.0%; ES – 9.7% and IR – 37.9% - Table 3) (Marquez et al., 2012; Reddy et al, 2013; Chamala et al., 2015). Of the alternative 3’ splice site events, 2,743 were of the NAGNAG type where two alternative 3’ splice sites are found 3 nt apart. Alternative NAGNAG 3’ splice sites can be of functional importance and are commonly found in human and plant genomes in coding sequences where they can add or remove a single amino acid and may be subject to regulation (Schindler et al., 2008; Busch and Hertel 2012; Shi et al., 2014).

**Table 3.** Frequencies of different alternative splicing events in BaRTv1.0

| Type of event | # | % |
| --- | --- | --- |
| Alternative 3' | 44,590 | 28.0% |
| Alternative 5' | 37,626 | 23.6% |
| Retained intron | 60,327 | 37.9% |
| Skipped exon | 15,387 | 9.7% |
| Mutually exclusive exons | 1,311 | 0.8% |
|  | 159,241 | 100.0% |

### Differential expression and differential alternative splicing in different barley organs/tissues

The major motivation for developing BaRTv1.0 was to exploit the fast, alignment-free transcript quantification software, Salmon, which requires an RTD to quantify transcript isoform abundances using k-mer indexing and counting (Patro et al., 2017). We used RNA seq data from three biological repeats of five organs/tissues of Morex to quantify transcripts with Salmon and BaRTv1.0-QUASI. Differential expression (DE) at both gene and transcript levels, differential AS (DAS) genes and differential transcript usage (DTU) were analysed using the recently developed 3D RNA-seq App (Calixto et al., 2018, 2019; Guo et al, personal communication). We removed poorly expressed transcripts from the dataset by stringent filtering (transcripts with ≥ 1 counts per million in at least 4 of the 15 samples were retained). A gene/transcript was significantly DE if it had an adjusted p-value of <0.01 and log_2_ fold change of ≥ 1. To identify significant DAS genes, consistency of expression changes (log_2_ fold change) between the gene and its transcripts was determined along with the change in splice ratio (Δ Percent Spliced – ΔPS). A DAS gene had at least one transcript which differed significantly from the gene and with an adjusted p-value of <0.01 and had at least a 0.1 (10%) change in ΔPS. Across the five organs and tissues, we detected expression of 60,807 transcripts from 25,940 genes. 20,972 genes were significantly DE across all tissues and 2,791 genes showed significant DAS (Figure 4A&D; Supplementary Table S6). The overlap between DE and DAS genes (those genes regulated by both transcription and AS) was 2,199 such that 592 genes were DAS-only and regulated only at the level of AS with no change in overall gene expression. We also identified 4,151 transcripts with significant DTU which underpins the differential AS. DTU transcripts behave differently from other transcripts of DAS genes and were identified by testing the expression changes of every transcript against the weighted average of all the other transcripts of the gene (Calixto et al., 2018). DTU transcripts differ significantly from the gene level and show a ΔPS of ≥0.1 with an adjusted p-value of <0.01. Pair-wise comparison of the number of up and down DE genes between each of the tissues showed that the two most related tissues (different developmental stages of inflorescence) had the fewest genes that were differentially expressed between them (ca. 700) but also had the highest number of DE genes when compared to other organs/tissues (ca. 14.5k between INF2 and NOD) (Figure 4B). There were ca. 10-fold fewer genes showing differential AS and pair-wise comparisons, which again showed that the two inflorescence tissues had the fewest numbers of DAS genes between them and INF2 compared to NOD, EMB and LEA had the highest numbers of DAS genes (Figure 4C). These results suggest that barley inflorescence transcriptomes differ substantially from shoot leaf, internode and embryos.

**Figure 4.**
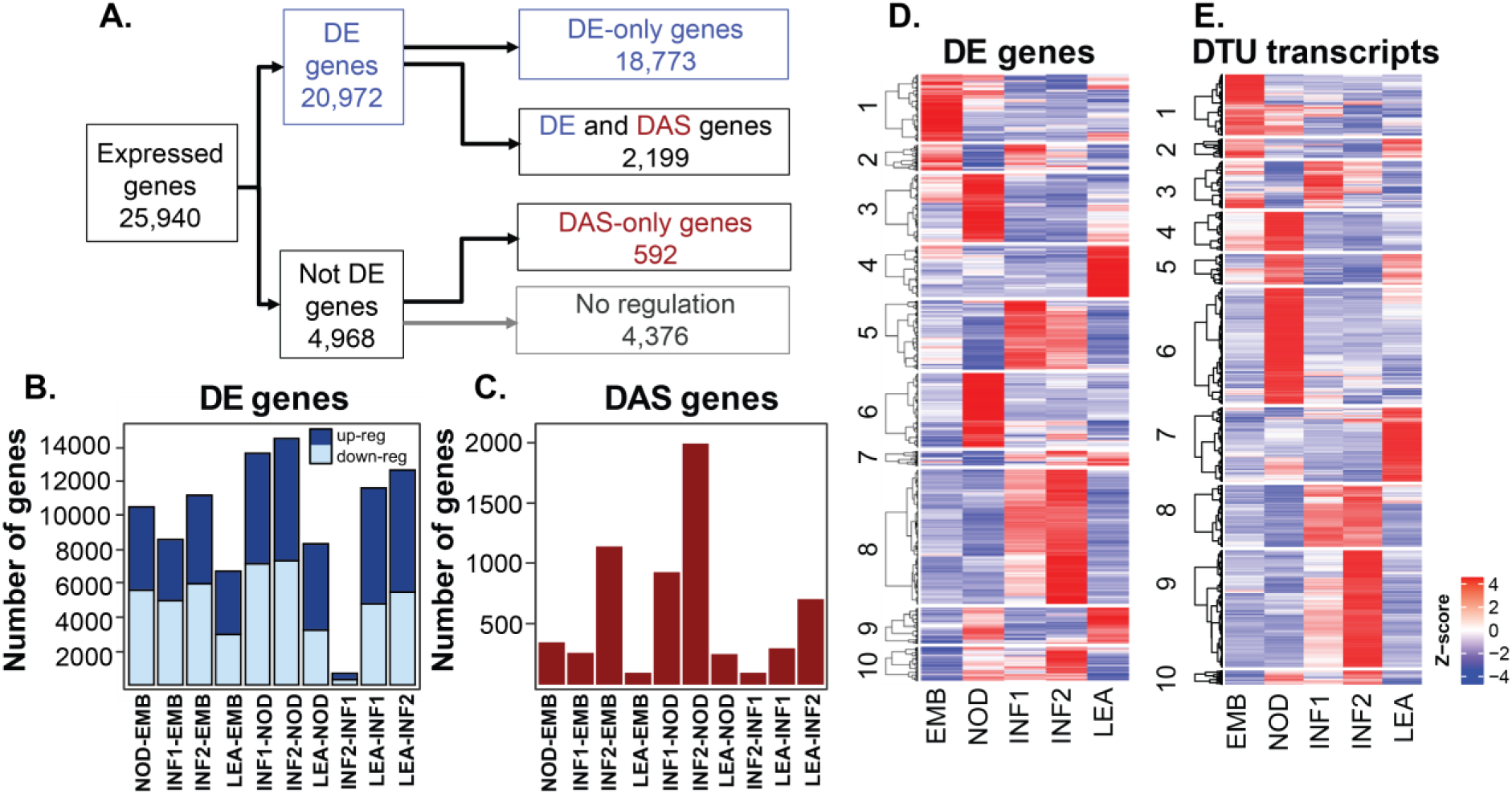
Differential gene and alternative splicing analysis in five barley organs. A. Numbers of expressed genes, differentially expressed genes (DE) and differential AS (DAS) across all 5 barley organs/tissues. B. Number of up- and down-regulated DE genes between pairs of different organs. Dark blue (up-regulated genes); light blue (down-regulated genes). C. Number of DAS genes between pairs of different organs. D. Heatmap and hierarchical clustering of 20,972 DE. E. Heatmap and hierarchical clustering of 2,768 DTU transcripts. The z-score scale in D and E represents mean-subtracted normalised log-transformed TPMs.

Hierarchical clustering of gene expression profiles of the 20,971 DE genes (DE-only and DE+DAS genes) across the organs/tissues identified clusters of genes that were co-ordinately and differentially expressed in each of the organs and tissues (Figure 4D). Cluster 1 (n=2,435) contained genes that were most highly expressed in the embryo, cluster 3 (n=2,477) and 6 (n=2,714) in the internode, cluster 5 (n=2,498) and 8 (n=4,906) in inflorescences and cluster 4 (n=1,880) and 9 (n=1,316) in leaf (Figure 4D; Supplementary Table S6). Hierarchical clustering also identified 2,768 transcripts differentially expressed DTU that showed some specificity of expression in each of the sampled tissues (Figure 4E; Supplementary Table S6). Cluster 1 (n=292) contains DTUs that are up-regulated in the embryo, Cluster 4, 5 and 6 (total n=885) in the internode and cluster 7 (total n=355) in shoot leaf. Cluster 3 (n=225) showed a cluster of DTU transcripts at the early stage of inflorescence development, cluster 8 (n=296) at both stages of inflorescence development and cluster 9 (n=559) at the later stage of inflorescence development. Thus, extensive differential gene and transcript expression and differential alternative splicing was revealed among the different samples using BaRTv1.0.

### Validation of differential AS from RNA-seq with HR RT-PCR and RNA-seq

To validate differential AS observed for individual genes among the different organs/tissues, we compared the RNA-seq quantifications of the 86 AS genes and 220 transcripts used in HR-RT-PCR. HR RT-PCR data showed over two-thirds of these transcripts had a significant differential AS (p = <0.001; >5% change) across the five samples (Supplementary Table S7). Given the RNA samples used in both the HR RT-PCR and RNA-seq was the same, we were able to directly compare differential AS observed at the individual gene level. For example, primer pairs Hv110 (HORVU5Hr1G027080; BART1_0-u34104) and Hv118 (HORVU1Hr1G078110; BART1_0-u5387) assay AS events that generate two alternative transcripts in BaRTv1.0. The AS transcripts are the result of alternative 5’ splice sites, 5 nt (Figure 5A) and 4 nt (Figure 5B) apart respectively. In each case selection of the distal 5’ splice sites produce the full-length CDS and use of the proximal 5’ splice site will result in a frame-shift and premature termination codons. Primer pair Hv173 (HORVU7Hr1G062930; BART1_0-u52907) assays alternative selection of two 3’ splice sites 33 nt apart (Figure 5C) and Hv217 (HORVU7Hr1G071060; BART1_0-u52404) assays retention of intron 1 (Figure 5D). Each of these examples show the pattern of AS across the tissues are essentially equivalent between HR RT-PCR and RNA-seq (Figure 5) and overall, we observed remarkable consistency. Thus, there is good agreement between the differential alternative splicing analysis from the RNA-seq data and the experimental verification with HR RT-PCR. These data provide strong support for the value of using BaRTv1.0 and BaRTv1.0-QUASI as reference datasets for accurate expression and AS analysis.

**Figure 5.**
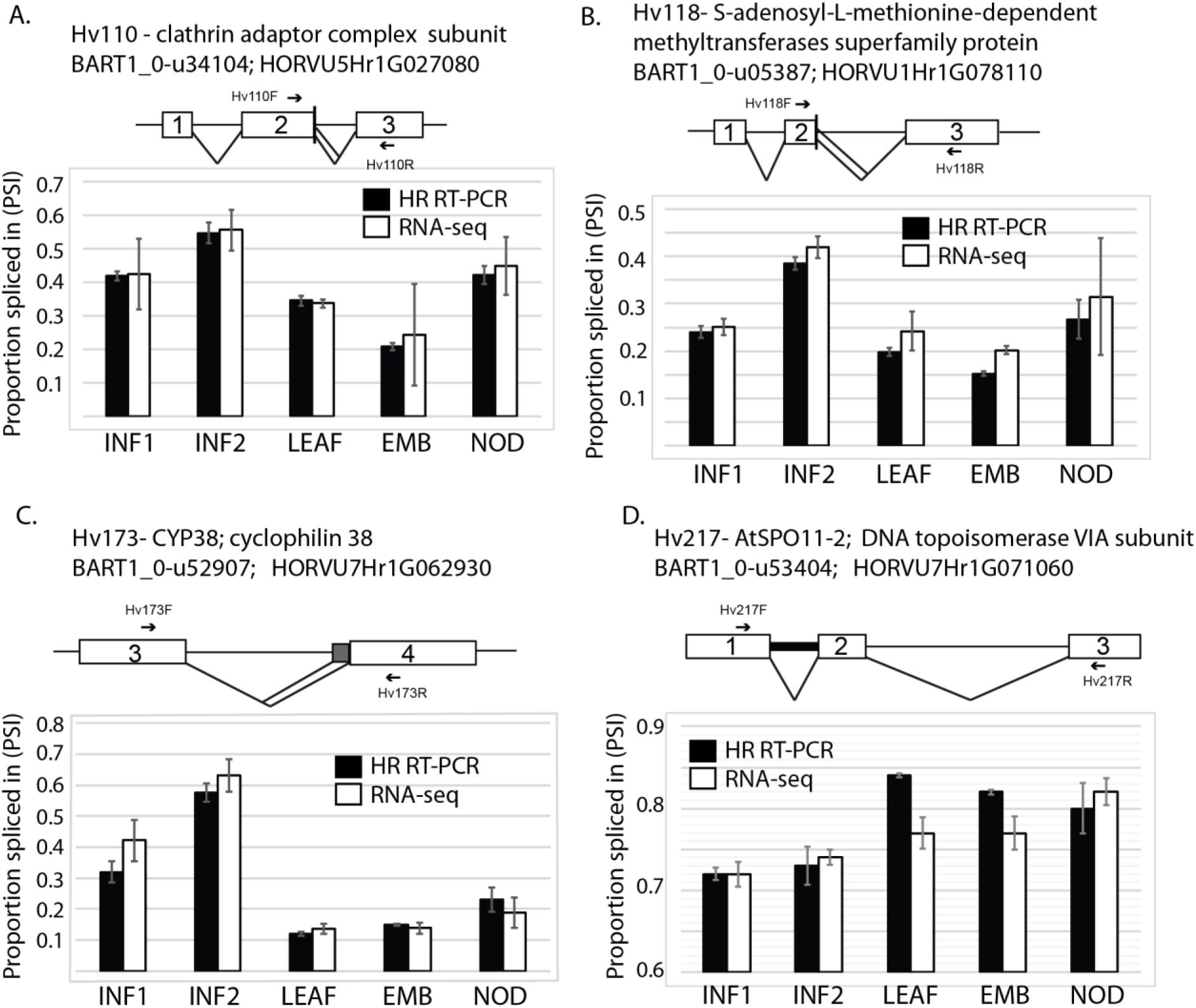
Comparison of alternative splicing in different barley tissues with HR RT-PCR and RNA-seq data. Splicing proportions of four different genes in 5 different barley tissues are presented. A. Hv110; HORVU5Hr1G027080, B. Hv118; HORVU1Hr1G078110, C. Hv173; HORVU7Hr1G062930, D. Hv217; HORVU7Hr1G071060. Schematic transcript/AS models are presented above histograms of PSIs derived from HR-RT-PCR (black) and RNA-seq (white) with standard error bars across three biological repeats. White boxes - exons, lines - introns; chevrons – splicing events; grey boxes region between alternative splice sites; thick intron line represents an intron retention.

### Complex patterns of AS

A principal aim of establishing BaRTv1.0 was to achieve higher accuracy of differential expression and AS analysis in barley RNA-seq datasets by improved transcript quantification. While the overall number of Morex transcripts in the HORVU collection (ca. 344k) was approximately halved in BaRTv1.0 (ca. 177k) (Table 1), some genes have multiple transcripts due to combinations of complex AS events. To examine the accuracy of such assembled transcripts we validated AS events in BART1_0-u51812, which codes for a WW domain-containing protein. BART1_0-u51812 contains 44 different transcript isoforms in the BaRTv1.0 dataset due to unique combinations of different AS events (Figure 6A). We analysed two regions that showed complex AS: between exons 2 and 3 and between exons 6 and 7 by HR RT-PCR (Figure 6). HR RT-PCR analysis identified fully spliced (FS), two alternative 5’ splice sites and retention of intron 2 as the main AS events between exons 2 and 3. In addition, four minor HR RT-PCR products were also identified and these were characterised as two further alternative 5’ splice sites and two alternative exons from the BaRTv1.0 transcripts (Figure 6B). Between exons 6 and 7, the main AS events are fully spliced, retention of intron 6, inclusion of an alternative exon and an alternative 5’ splice site (Figure 6C). HR RT-PCR across exons 6-7 (primer pair Hv79 in exons 6 and 8) accurately identified these AS events (Figure 6C). These AS events were also quantified using transcript abundances from the RNA-seq data using BaRTv1.0_QUASI and showed good agreement with the HR RT-PCR results with Pearson correlations of 0.92 for the Hv78 regions and 0.73 for the Hv79 region. These examples support the accuracy of alternative transcripts found in BaRTv1.0 and that transcripts with complex combinations of AS events can be quantified in RNA-seq data with confidence.

**Figure 6.**
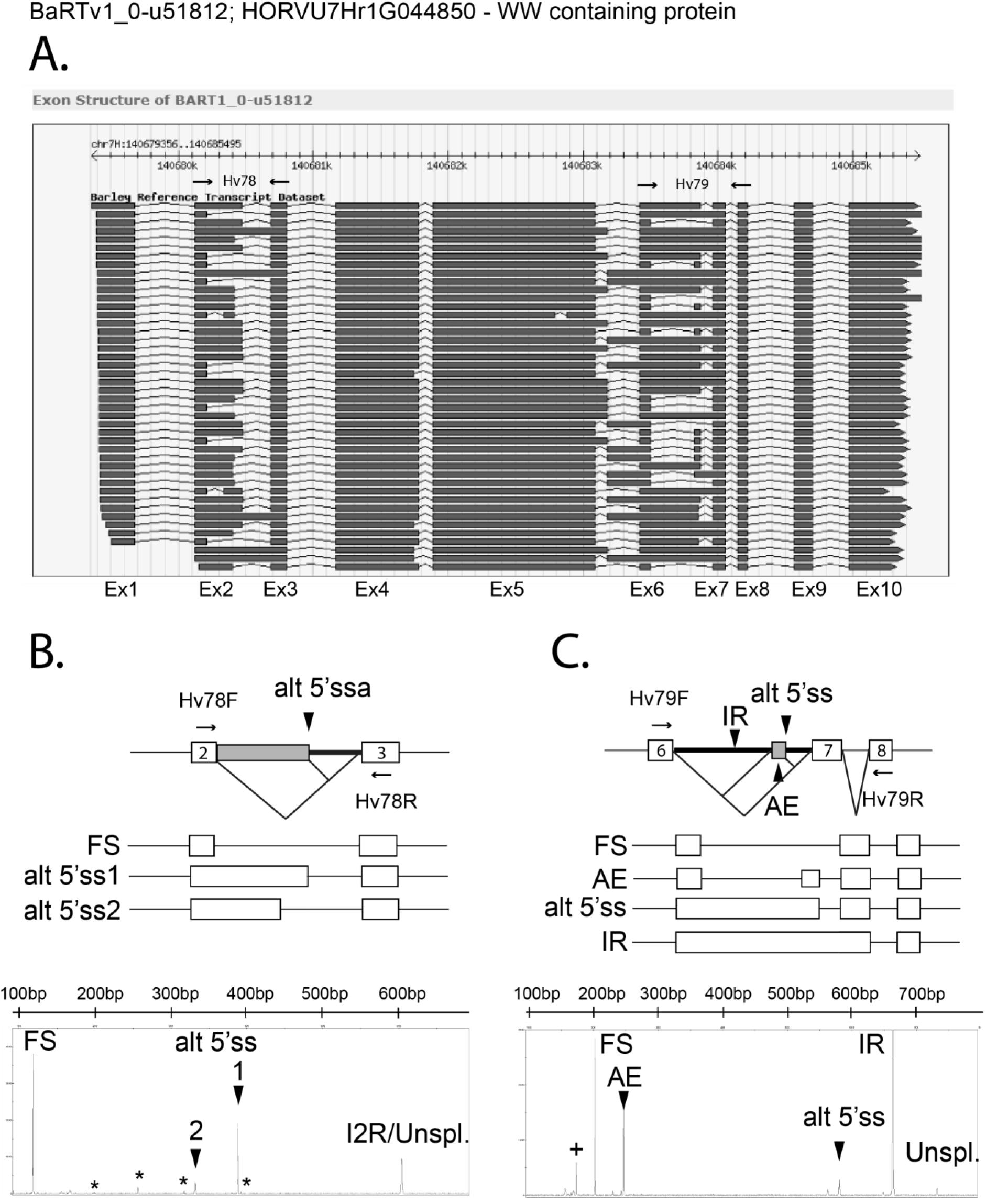
Alternative splicing in a WW domain containing protein gene (BART1_0-u51812). A. BART1_0-u51812 transcript models represented in the BaRTv1.0 database. B. AS events involving intron 2 validated by HR-RT-PCR. C. AS events between exon 6 and 8 validated by HR-RT-PCR. Electropherogram output from the ABI3730 shows the HR RT-PCR products (x-axis RT-PCR products (bp); y-axis relative fluorescence units). The products expected from RNA-seq are indicated as FS – Fully spliced, AE - Alternative exon, Alt 5’ss - Alternative 5’ splice site, IR-intron retention and Unspl.-Unspliced. * in B. indicates minor alternative transcripts identified in HR RT-PCR and in RNA-seq. + in C. indicates an uncharacterised alternative transcript identified in HR RT-PCR.

## Discussion

Comprehensive reference transcript datasets are required for accurate quantification of transcripts for expression analysis using RNA-seq. Quantification at the transcript level improves gene level expression estimates and allows robust and routine analysis of alternative splicing. Here we describe the BaRTv1.0 transcript dataset or transcriptome for barley, produced by merging and filtering transcripts assembled from extensive RNA-seq data and its utility in differential expression and differential alternative splicing. The transcripts were assembled against cv. Morex and this reference transcript dataset is therefore a Morex assembly. BaRTv1.0 achieves a balance between maximising transcript diversity – all 177,240 transcripts have at least one unique splice site with strong junction support – and reducing the numbers of mis-assembled transcripts, transcript fragments and redundant transcripts. This barley transcript dataset represents the first stage of an evolving resource which will continue to improve and expand as more complete barley genomes are released and by incorporation of new Illumina short read data along with single molecule sequencing (Pacific Biosciences or Oxford Nanopore Technology) datasets when they become available. Long-read data will confirm transcript features proposed by the short-read assemblies by defining the exact combinations of different AS events and 5’ and 3’ ends and may identify rare transcripts. The transcript and splice junction data generated here will be valuable in improving the barley genome annotation. Finally, the BaRTv1.0 transcript dataset will enable accurate gene and transcript level expression and AS analysis increasing our understanding of the full impact of AS and how transcriptional and AS regulation of expression interact to determine barley development, responses to environment and ultimately important crop phenotypes such as yield, disease resistance and stress tolerance.

BaRTv1.0 represents 60,444 genes, which is considerably fewer than the 81,683 genes reported in the current barley genome (Mascher et al., 2017) where residual gene fragmentation has likely inflated the number of annotated genes. This number of genes is still higher than expected and may decrease with further improvements in genome annotation. However, the arrangement of BaRTv1.0 transcripts have identified mis-annotated chimeric genes in the barley reference genome, helping to improve gene resolution. BaRTv1.0 was established using RNA-seq data containing approximately 19 billion reads from a range of different biological samples (organs, tissues, treatments and genotypes) and was assembled initially against the Morex genome. The sequence depth and rigorous filtering and validation allowed us to establish a diverse set of high-quality, robust and experimentally supported transcripts.

A key function of the BaRTv1.0 transcript dataset is improved accuracy of transcript abundance. Variation in the 5’ and 3’ ends of transcripts of the same gene was shown previously to affect transcript quantification in Arabidopsis (Zhang et al., 2017a) and similar results for 3’ end variation have been found in human RNA-seq analysis (Soneson et al., 2019). Extending the sequences of shorter transcripts with genomic sequences such that all transcripts of a gene had the same 5’ and 3’ ends improved the accuracy of transcript quantification compared to experimental data (Zhang et al., 2017a). We also found an improvement in the quantification of transcripts and splicing proportions by applying the same approach to produce the BaRTv1.0-QUASI version, specifically for quantification of alternatively spliced isoforms (Table 1). The continued development of reference transcript datasets for other lines and cultivars will be essential for accurate gene expression and AS analysis. One significant application will be to enable genome-wide association studies using gene expression data to identify eQTLs and transcript abundance/splicing ratios to identify splicing QTLs (Thatcher et al., 2014).

To demonstrate the value of the new RTD for gene expression studies and AS analysis, we used BaRTv1.0-QUASI to quantify transcripts in the five developmental organs and tissues RNA-seq datasets that we had used previously for HR RT-PCR optimisation and validation. We observed extensive differences in gene expression and AS among the five divergent samples. Clustered co-expression patterns clearly showed that the different organs and tissues have distinct transcriptomes reflecting major differences in both transcription and AS, as recently demonstrated in the cold response in Arabidopsis (Calixto et al., 2018). The abundance of individual BaRT transcripts in these five organs/tissues, and in the eleven other organs and tissues used to annotate the barley genome (Mascher et al., 2017) are displayed in a barley reference transcript database website https://ics.hutton.ac.uk/barleyrtd/index.html.

Barley is adapted to a wide range of environments and is grown for many purposes. As a result, different cultivars/genotypes will have unique transcriptome profiles that will respond differently to varying developmental or environmental conditions and challenges. BaRTv1.0 enables rapid and robust analysis of gene expression and AS in a wide range of experimental scenarios. BaRTv1.0 is based on cv. Morex but used RNA-seq data from a wide-range of cultivars and lines. We anticipate significant and incremental improvements in subsequent BaRT iterations by adding new short and long-read RNA-seq datasets but understand the need to capture the diversity of different transcripts which will occur among different cultivars and landraces. Sequence variation among different lines will generate quantitative variation in expression and alternative splicing (Gan et al., 2011). Therefore, using the methods presented here, RTDs for other widely used cultivars can be generated. For example, construction of RTDs for Golden Promise (used for genetic transformation studies) (Mrízová et al., 2014), Bowman (the background cultivar for a collection of near isogenic lines) (Dahleen et al., 2005) and Barke (a cultivar more relevant to modern European cultivated barley) (Pham et al., 2019) would all have specific utility. Ultimately, transcript data from a wide range of genotypes will stimulate the move towards the development of a reference pan-transcriptome to parallel the generation of the barley pan-genome sequence.

## Conclusions

A comprehensive, non-redundant barley reference transcript dataset called BaRTv1.0 has been generated, which enables fast, precise transcript abundances. Downstream analysis of transcript abundances in five barley organs/tissues, identified significant differential expression of many genes and transcripts. BaRTv1.0 is part of a unique pipeline that facilitates the robust routine analysis of barley gene expression and AS. The reference transcripts have broader opportunities to develop unique expression markers, support proteomic resources for barley and enable transcript/co-expression/regulatory networks. The pipeline developed here has relevance to the development of other crop reference transcript datasets.

## Materials and Methods

An experimental and bioinformatics workflow showing the assembly, filtering and validation approach taken is shown in Figure 1.

### Selected RNA-seq datasets and data processing

A total of 11 large RNA-seq datasets consisting of 808 samples including replicates, were selected to assemble a barley transcriptome (Supplementary Table S1). Eight publicly available datasets were downloaded from NCBI - Sequence Read Archive database (https://www.ncbi.nlm.nih.gov/sra/) and the 3 remaining datasets are currently unpublished. All datasets were produced using Illumina platforms and were selected based on being the most recent datasets with the longest read length available (mostly >90 bp and paired-end reads) with a quality of q >=20. All raw data were processed using Trimmomatic-0.30 (Bolger et al., 2014) using default settings to preserve a minimum Phred score of Q20 over 60 bp. One of the samples (NOD1) was over-represented with respect to read numbers due to a repeat run being necessary, and was therefore subsampled to 60 million reads. Read quality before and after trimming was performed using FastQC (fastqc_v0.11.5) (https://www.bioinformatics.babraham.ac.uk/projects/fastqc/).

### Transcriptome assembly

#### Alignment

Transcript assembly was performed using a data pipeline that initially used STAR (version 2.5; Dobin et al., 2013) to align reads from each of the 808 samples individually to the latest barley cv. Morex reference genome (version 160404_barley_pseudomolecules_parts_masked /Hv_IBSC_PGSB_v2) (Mascher et al., 2017). Many alignment programmes use a two-step approach to identify exon junctions and then use the junctions to guide the final alignment (Engström et al., 2013). A three-step STAR alignment approach was developed to improve alignment accuracy and identification of splice junctions and to take into consideration the sequence variation in reads from different cultivars and lines used and to capture splice junctions from tissue/conditions samples where the amount of material or sequencing depth were limited or where genotypes were represented by small numbers of samples. In the first pass, reads were mapped to the genome allowing a single mismatch and only those with an overhang minimum of 10 bp on each side of the splice junction were taken forward. This step identified 1,057,650 splice junctions, many of which were supported by only a single read. These splice junctions were filtered based on their read support requiring 5 or more uniquely mapped reads or 10 or more reads if multi-mapped reads were present; the remaining 206,688 splice junctions were used as annotation for the second pass. In the second pass the alignment was relaxed to allow 2 mismatches in the splice junction region with an overhang minimum of 7 bp. This step identified 1,088,440 splice junctions and these were further filtered to select splice junctions on the basis of three sets of criteria: a) splice junctions with 3 or more uniquely mapped reads (5 or more reads if multi-mapped reads are present) in at least 2 samples; b) splice junctions with 2 or more uniquely mapped reads in at least 5 samples or c) splice junctions supported by 1 or more uniquely mapped reads in at least 10 samples and allowing for 2% mismatches in the alignment of reads outside the splice junction. In the final pass, the 323,619 filtered splice junctions from the previous step were used as annotation and no new splice junctions were allowed. In this step, the read mismatch rate was relaxed to 3% to allow more reads to map. In all three passes, only canonical splice junctions (GT..AG, GC..AG and AT..AC) and concordant alignments were kept.

#### Transcript assembly

After STAR alignment, each sample was run individually using StringTie (version 1.3.3b) (Pertea et al., 2015). Different combinations of StringTie parameters were extensively tested and the parameters that produced the best assembly were retained (see Results). Evaluation of each assembly was performed based on comparison with HR RT-PCR data consisting of 86 genes and 220 alternatively spliced RT-PCR products (see Results). To evaluate the completeness of the transcripts assembled, 22,651 Haruna nijo fl-cDNAs (Matsumoto et al., 2011) were aligned using BLASTn (blastn, version ncbi-blast-2.2.28+; Altschul SF et al, 1990) to each RNA-seq transcriptome assembly generated. All fl-cDNAs with ≥ 90% coverage and ≥ 90% identity were identified and the total number was considered a measure of completeness. Optimal StringTie parameters were coverage (-c 2.5); gap between readings triggering a new bundle (-g 50); isoform fraction was set at -f 0, gene abundance estimation was set as output (-A), minimum anchor length for junctions 5 (-a); minimum junction coverage 0.1 (-j) and fraction of bundle allowed to be covered by multi-hit reads 1 (-M).

#### Removal of low abundance transcripts

Salmon is a software tool that utilises a defined set of reference sequences to perform a rapid, alignment-free estimation of isoform abundances by using k-mer indexing and counting. It uses an accelerated expectation-maximization algorithm for quantifying isoform abundance, which is given in transcripts per million (TPM). All 808 individual StringTie assemblies were merged with StringTie-merge, after all 808 read samples were aligned to the merged reference transcriptome with Salmon (version Salmon-0.8.2) (Patro et al, 2017) to obtain transcript quantification. All transcripts that were expressed at less than 0.3 TPM, across all samples, were filtered out.

#### Assembly merge

All 808 assembly predictions from StringTie were merged using StringTie-merge to create a unique consensus assembly version. A minimum isoform fraction of 0 (-f) and a minimum input transcript TPM of 0.1 (-T) was used in StringTie-merge. The consensus transcriptome, after filtering out the transcripts less than 0.3 TPM, was further merged (gtf format) with the 22,651 Haruna nijo (HN) fl cDNAs (Matsumoto et al., 2011). The HN fl cDNAs were mapped to the barley cv. Morex genome with the GMAP tool (version 2017-10-30) (Wu & Watanabe, 2005). Finally, we used TransDecoder (version 5.3.0) (Haas et al., 2013) and BLASTp to identify and filter out all transcripts equal to or less than 300 bp (8,831 transcripts) with less than 70% of coverage and identity protein homology with the protein datasets from 3 reference Poaceae species – *Oriza sativa* (v7_JGI), *Brachypodium distachyon* (Bd21-3 v1.1) and *Sorghum bicolor* (v3.1.1) (https://genome.jgi.doe.gov/portal/) (Supplementary Figure 4) to establish BaRTv1.0.

### Alternative splicing analysis

The newly created non-redundant BaRTv1.0 consensus transcriptome was further refined to allow accurate quantification of AS as described previously, to create a separate dataset specifically for quantification of AS isoforms (BaRTv1.0 – QUASI) (Zhang et al., 2017a). All transcripts with shorter 5’ and 3’ UTR regions were padded out to the 5’ and 3’ ends of the longest transcript of that gene using the cv. Morex genome.

### Comparing HR RT-PCR and RNA-seq alternative splicing proportions

To assess the accuracy of BaRTv1.0 to detect changes in AS in the RNA-seq data, we compared the splicing proportions for AS events from HR RT-PCR with those calculated from the RNA-seq data using the HORVU transcript set, BaRTv1.0 and BaRTv1.0-QUASI as transcript references. To establish the correlations, a number of considerations were required. First, HR RT-PCR data reports exclusively on the events that occur within a gene bordered by the primers used for the analysis. The RNA-seq data reports on individual transcripts that may contain multiple AS events or have an alternative transcript start and/or stop. For this reason, multiple RNA-seq transcripts may represent the same AS product that is detected by HR RT-PCR. We therefore developed a method (https://github.com/PauloFlores/RNA-Seq-validation) that determined the size of the expected PCR product by aligning the primer pairs against each RNA-seq transcript and determining the predicted length that PCR would produce. The TPM values of all transcripts that produce the same AS PCR product were added together to give a combined RNA-seq value for that PCR product. The proportions of the different AS products for both HR-RT-PCR and RNA-seq were then subsequently calculated and correlated.

Firstly, the method mapped the HR RT-PCR primers to the transcriptome using BLAST (blastn-short command; version ncbi-blast-2.2.28+; Altschul et al, 1990). All transcripts with perfect identity and coverage for both reverse and forward primers at one gene transcript location were selected (http://ics.hutton.ac.uk/barleyrtd/primer_list.html). Secondly, the distance was calculated between the pairs of primers for each selected transcript, and thirdly, transcripts with equal product length associated with the same pair of primers were clustered together. Fourthly, five reference samples from the sample dataset, each with 3 biological replicates to give 15 datasets (IBSC, 2012) were individually quantified by Salmon (version Salmon-0.8.2; Patro et al, 2017). The five reference samples consisted of 4-day old embryos dissected from germinating grains (EMB), young developing inflorescences (5 mm) (INF1), developing inflorescences (1-1.5 cm) (INF2), developing tillers at 6 leaf stage, third internode (NOD) and shoots from seedlings (LEA). The levels of expression (in TPM) from Salmon were summed for transcripts with the same RT-PCR product lengths. For each pair of primers and allowing for a difference of ±6 bp (to allow for inaccuracies in HR RT-PCR size calling), products of the same length between HR RT-PCR and RNA-seq were identified. Finally, based on the calculated values of RNA-seq levels of expression and the calculated values of HR RT-PCR for each RT-PCR product, the proportions of the alternative transcripts were calculated (transcriptTPM/Summed_transcriptsTPM compared to RT-PCR productRFU/Summed_RT-PCRproductRFU) and the Pearson’s and Spearman’s correlation co-efficient was calculated (see Supplementary Figure 5 for a pipeline summary).

### Percent spliced in values and identification of alternative splicing type

SUPPA version 2.3 (Alamancos et al., 2015) determined AS events and calculated the relative inclusion values of AS events. Outputs from Salmon were fed into SUPPA to quantify AS events across the tissue sample datasets and generate percentage spliced in (PSI) values.

### Generation of the BaRTv1.0 database

A database and website front-end were constructed to allow easy access to BaRTv1.0 transcripts and expression analyses using the LAMP configuration (Linux, Apache, mySQL, and Perl). Additional annotation was added to the transcripts by homology searching against the predicted peptides from rice (rice pseudo-peptides v 6.0; Ouyang et al., 2007) and from Arabidopsis thaliana (TAIR pseudo-peptides v 10, The Arabidopsis Information Resource) using BLASTX at an e-value cutoff of less than 1e-50 (Altschul et al., 1990). The website https://ics.hutton.ac.uk/barleyrtd/index.html allows users to interrogate data through an entry point via three methods: (i) a BLAST search of the reference barley assembly or the predicted transcripts; (ii) a keyword search of the derived rice and Arabidopsis thaliana BLAST annotation, and; (iii) a direct string search using the transcript, gene, or contig identifiers. To distinguish this new set of predicted genes and transcripts from previously published ‘MLOC_’ and HORVU identifiers, they have subsequently been assigned a prefix of ‘BART1_0-u00000’ for the unpadded or ‘BART1_0-p00000’ for the padded QUASI version, with 00000 representing the individual transcript number.

The RNA-seq TPM values for the developmental stages of barley (Morex cultivar) (Mascher et al 2017) at the replicate and stage are shown in both graphic and tabular formats for each gene. The exon structures of the transcripts for each gene are shown in graphical form, and links to the transcripts themselves provides access to the transcript sequences in FASTA format. Each transcript has also been compared to the published set of predicted genes (HORVUs) in order to provide backwards compatibility.

### High resolution RT-PCR

The RNA from five of sixteen developmental stages of barley cv.Morex was used for HR RT-PCR validation (Mascher et al., 2017). This consisted of three biological replicates of leaf tissue (LEA) sampled from seedlings at 17 days after planting (dap); the third stem internode (NOD) dissected at 42 dap; whole developing inflorescence tissue sampled at 30 dap (INF1) and 50 dap (INF2) and embryonic tissue (including mesocotyl and seminal roots; EMB) dissected after 4 days. High resolution RT-PCR was performed essentially as described previously (Simpson et al., 2008). A panel of 86 primer pairs covering 220 RT-PCR products (Supplementary Table S3), were designed to barley genes that showed evidence of AS and more than 100 RNA-seq reads for each primer pair to support transcription, with the exception of 14 primer pairs numbered between primers #14 and 51. These primers were designed to genes already under study and consisted of splicing factor genes, clock response genes and Rubisco activase (Supplementary Table S3). Primers were designed to amplify products between 100 and 700 bp to capture the different splicing events. The 5’ upstream primer was 5’ labelled with 6-Carboxyfluorescein (6-FAM). Total RNA (5 μg) was used for first-strand cDNA synthesis by reverse transcription with oligo(dT)18 using Ready-To-Go You-Prime First-Strand Beads (GE Healthcare) in a final volume of 20 μL. RT-PCR was performed as described (Simpson et al., 2008) and the resultant RT-PCR products representing AS transcripts were detected on an ABI3730 DNA Analyzer (Thermo Fisher Scientific) along with GeneScan 500 LIZ size standard (Applied Biosystems). RT-PCR products were accurately sized and peak areas calculated (Relative Fluorescence Units – RFUs) using GeneMapper (ABI) software.

### Statistical analysis

#### HR RT-PCR ANOVA

Pairwise significance of the variation between the developmental tissues was assessed by analysis of variance (ANOVA). Each peak of each primer was analysed separately with three replicate values for each treatment combination. Response was measured as the percentage contribution of a particular isoform to the total transcripts measured, and ANOVA was carried out after an angular (arcsin) transformation was used to transform values from [0, 1] to [-π/2, + π/2] to give the data a normal distribution (Sokal and Rohlf, 1995). Fisher’s Least Significant Difference (LSD) test was performed for the pairwise comparisons between the different tissues tested at a p- value <0.001. In the subsequent analysis, we focussed on those transcripts which showed a significant increase or decrease with a 5% difference between the means of the different plant tissues. This level of difference was selected because we previously determined that when comparing variation in technical reps in the AS RT–PCR system, the majority of transcripts showed a standard error of the mean of <3% (Simpson et al., 2008; Kim et al., 2009).

## Declarations

## Data Access

BaRTv1.0 and BaRTv1.0 – QUASI are available as .fasta and .GFF files and can be downloaded from https://ics.hutton.ac.uk/barleyrtd/downloads.html.

## Author Contributions

PR-F, MB, CS, C-DM, and RW conceived and designed the experiment and AS analysis. PR-F and C-DM performed the statistical and correlation analyses. PEH and JM prepared the RNA samples for the HR RT-PCR analysis and co-ordinated the sequencing. JF, JWSB, CS RW, AB, MS, CH, JK, SM, MCC and MZ generated RNA and libraries for RNA-seq. PR-F and MB assembled the RNA-seq data. JF, GS and CS identified the AS genes and performed the HR RT-PCR screening and analysis of the data. LM established the searchable database. PR-F, C-DM, JWSB, RZ, WG and CS performed the detailed analysis of the RNA-seq and HR RT-PCR data. CS, PR-F, RW and JWSB wrote the paper.

## Acknowledgements and Funding

This research was supported by Scottish Government Rural and Environment Science and Analytical Services division (RESAS) and funding from the Biotechnology and Biological Sciences Research Council (BBSRC) (BB/I00663X/1: A draft sequence of the barley genome, to RW). RNA-seq datasets were produced through funding to CH through the BBSRC Sustainable Bioenergy Centre initiative grant no. BB/G016232/1, BBSRC Response Mode Grant No. BB/K01613X/1 to SM and RW and ERC project 669182 ‘SHUFFLE’ to RW. We further acknowledge those involved with the ERC project 669182 ‘SHUFFLE’ project in generating samples

