## Supplementary figures for "BaRTv1.0: an improved barley reference transcript dataset to determine accurate changes in the barley transcriptome using RNA-seq"

### Slide 1
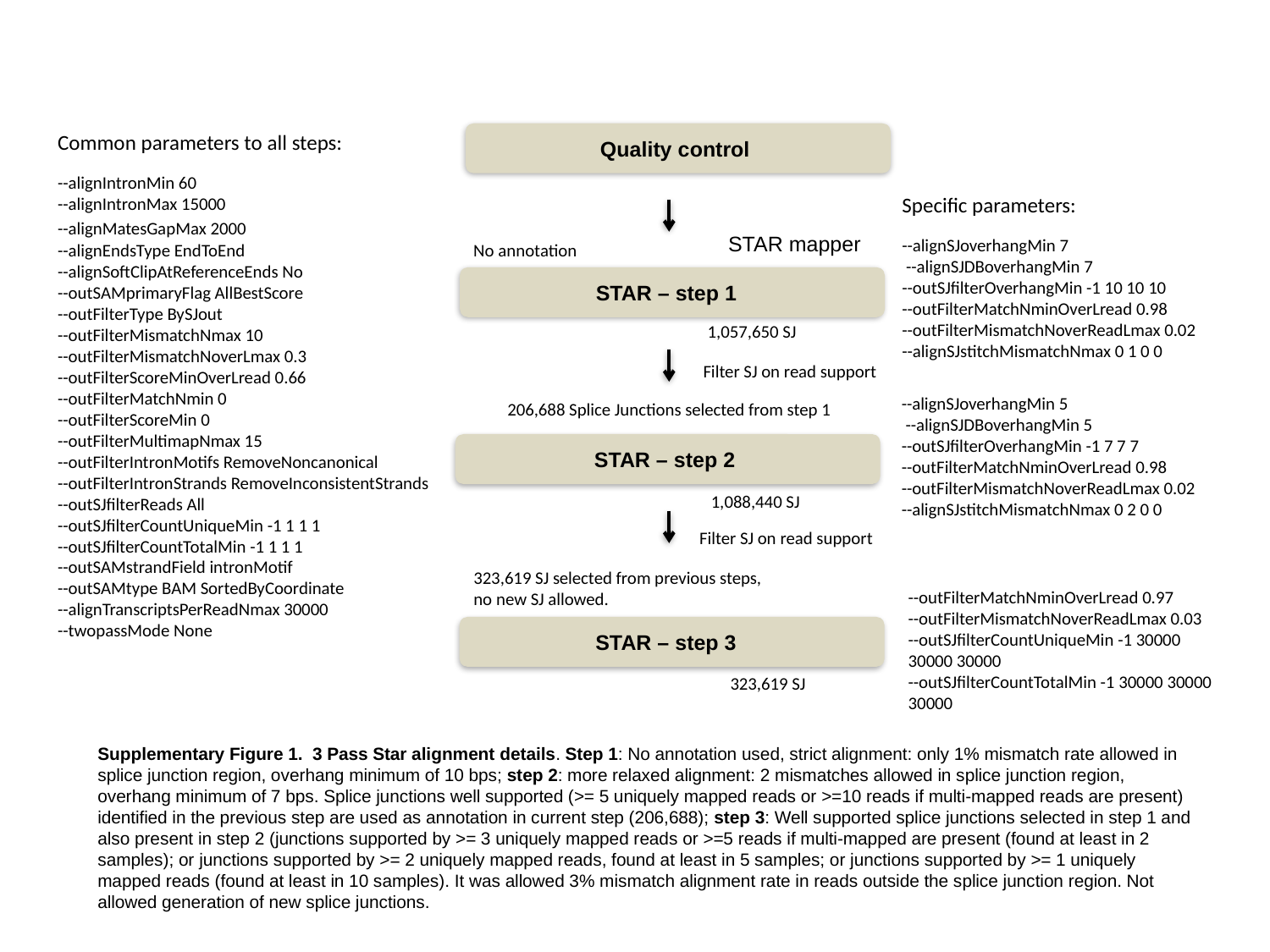

Common parameters to all steps:
--alignIntronMin 60
--alignIntronMax 15000
--alignMatesGapMax 2000
--alignEndsType EndToEnd
--alignSoftClipAtReferenceEnds No
--outSAMprimaryFlag AllBestScore
--outFilterType BySJout
--outFilterMismatchNmax 10
--outFilterMismatchNoverLmax 0.3
--outFilterScoreMinOverLread 0.66
--outFilterMatchNmin 0
--outFilterScoreMin 0
--outFilterMultimapNmax 15
--outFilterIntronMotifs RemoveNoncanonical
--outFilterIntronStrands RemoveInconsistentStrands
--outSJfilterReads All
--outSJfilterCountUniqueMin -1 1 1 1
--outSJfilterCountTotalMin -1 1 1 1
--outSAMstrandField intronMotif
--outSAMtype BAM SortedByCoordinate
--alignTranscriptsPerReadNmax 30000
--twopassMode None
Quality control
Specific parameters:
--alignSJoverhangMin 7
 --alignSJDBoverhangMin 7
--outSJfilterOverhangMin -1 10 10 10
--outFilterMatchNminOverLread 0.98
--outFilterMismatchNoverReadLmax 0.02
--alignSJstitchMismatchNmax 0 1 0 0
STAR mapper
No annotation
STAR – step 1
1,057,650 SJ
Filter SJ on read support
--alignSJoverhangMin 5
 --alignSJDBoverhangMin 5
--outSJfilterOverhangMin -1 7 7 7
--outFilterMatchNminOverLread 0.98
--outFilterMismatchNoverReadLmax 0.02
--alignSJstitchMismatchNmax 0 2 0 0
206,688 Splice Junctions selected from step 1
STAR – step 2
1,088,440 SJ
Filter SJ on read support
323,619 SJ selected from previous steps,
no new SJ allowed.
--outFilterMatchNminOverLread 0.97
--outFilterMismatchNoverReadLmax 0.03
--outSJfilterCountUniqueMin -1 30000 30000 30000
--outSJfilterCountTotalMin -1 30000 30000 30000
STAR – step 3
323,619 SJ
Supplementary Figure 1. 3 Pass Star alignment details. Step 1: No annotation used, strict alignment: only 1% mismatch rate allowed in splice junction region, overhang minimum of 10 bps; step 2: more relaxed alignment: 2 mismatches allowed in splice junction region, overhang minimum of 7 bps. Splice junctions well supported (>= 5 uniquely mapped reads or >=10 reads if multi-mapped reads are present) identified in the previous step are used as annotation in current step (206,688); step 3: Well supported splice junctions selected in step 1 and also present in step 2 (junctions supported by >= 3 uniquely mapped reads or >=5 reads if multi-mapped are present (found at least in 2 samples); or junctions supported by >= 2 uniquely mapped reads, found at least in 5 samples; or junctions supported by >= 1 uniquely mapped reads (found at least in 10 samples). It was allowed 3% mismatch alignment rate in reads outside the splice junction region. Not allowed generation of new splice junctions.

### Slide 2
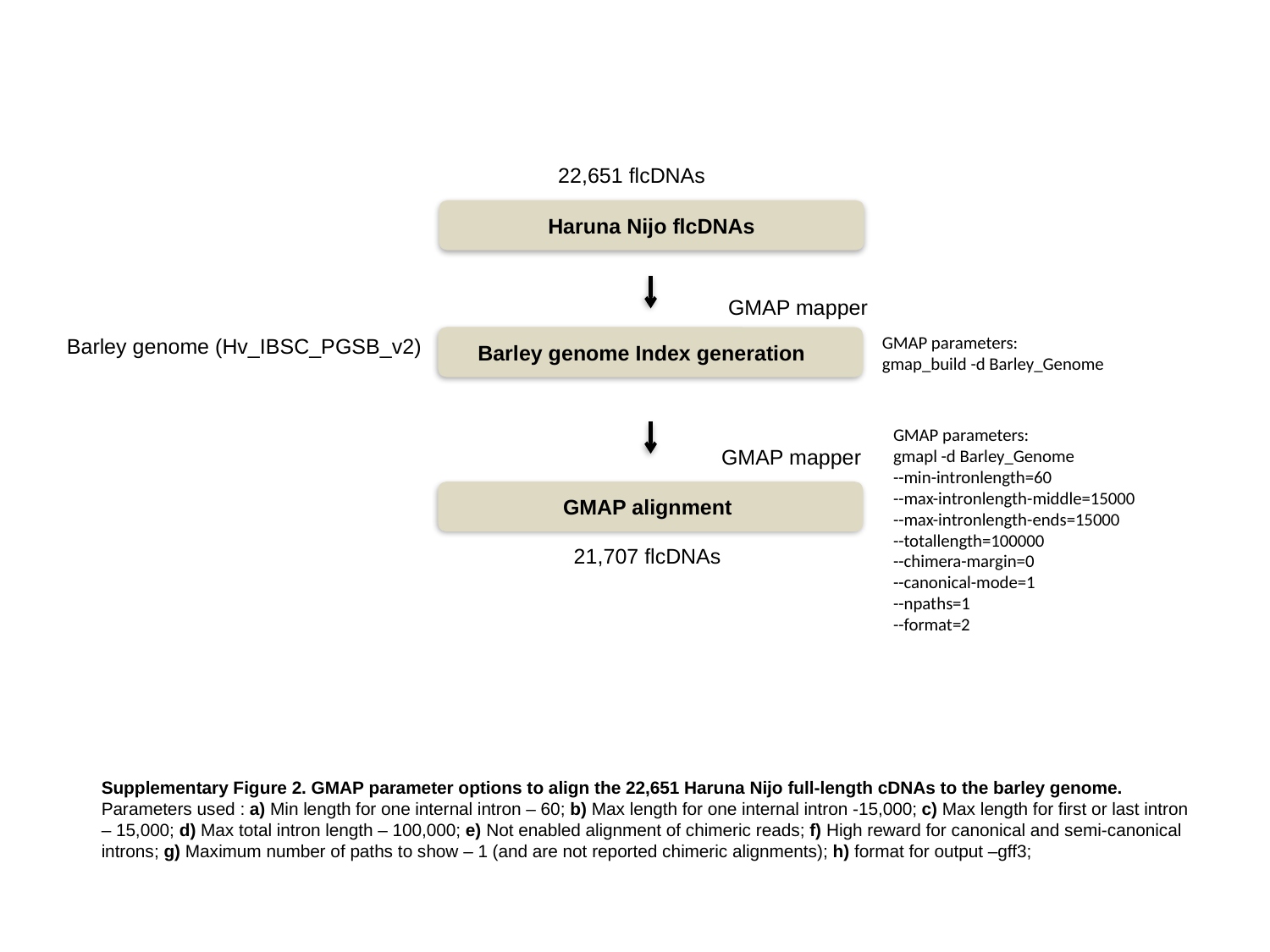

22,651 flcDNAs
Haruna Nijo flcDNAs
GMAP mapper
GMAP parameters:
gmap_build -d Barley_Genome
Barley genome (Hv_IBSC_PGSB_v2)
Barley genome Index generation
GMAP parameters:
gmapl -d Barley_Genome
--min-intronlength=60
--max-intronlength-middle=15000
--max-intronlength-ends=15000
--totallength=100000
--chimera-margin=0
--canonical-mode=1
--npaths=1
--format=2
GMAP mapper
GMAP alignment
21,707 flcDNAs
Supplementary Figure 2. GMAP parameter options to align the 22,651 Haruna Nijo full-length cDNAs to the barley genome. Parameters used : a) Min length for one internal intron – 60; b) Max length for one internal intron -15,000; c) Max length for first or last intron – 15,000; d) Max total intron length – 100,000; e) Not enabled alignment of chimeric reads; f) High reward for canonical and semi-canonical introns; g) Maximum number of paths to show – 1 (and are not reported chimeric alignments); h) format for output –gff3;

### Slide 3
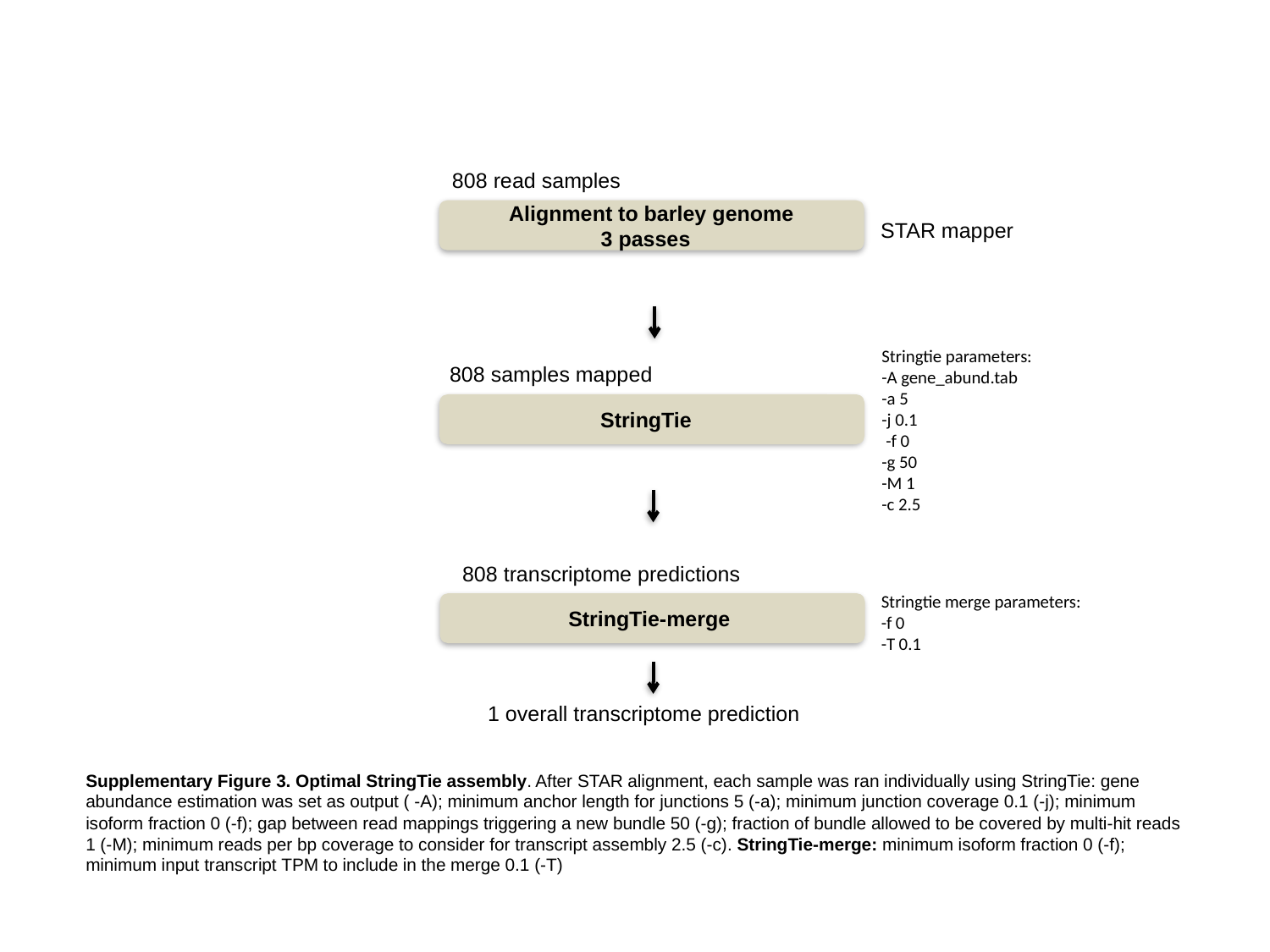

808 read samples
Alignment to barley genome
3 passes
STAR mapper
Stringtie parameters:
-A gene_abund.tab
-a 5
-j 0.1
 -f 0
-g 50
-M 1
-c 2.5
808 samples mapped
StringTie
808 transcriptome predictions
Stringtie merge parameters:
-f 0
-T 0.1
StringTie-merge
1 overall transcriptome prediction
Supplementary Figure 3. Optimal StringTie assembly. After STAR alignment, each sample was ran individually using StringTie: gene abundance estimation was set as output ( -A); minimum anchor length for junctions 5 (-a); minimum junction coverage 0.1 (-j); minimum isoform fraction 0 (-f); gap between read mappings triggering a new bundle 50 (-g); fraction of bundle allowed to be covered by multi-hit reads 1 (-M); minimum reads per bp coverage to consider for transcript assembly 2.5 (-c). StringTie-merge: minimum isoform fraction 0 (-f); minimum input transcript TPM to include in the merge 0.1 (-T)

### Slide 4
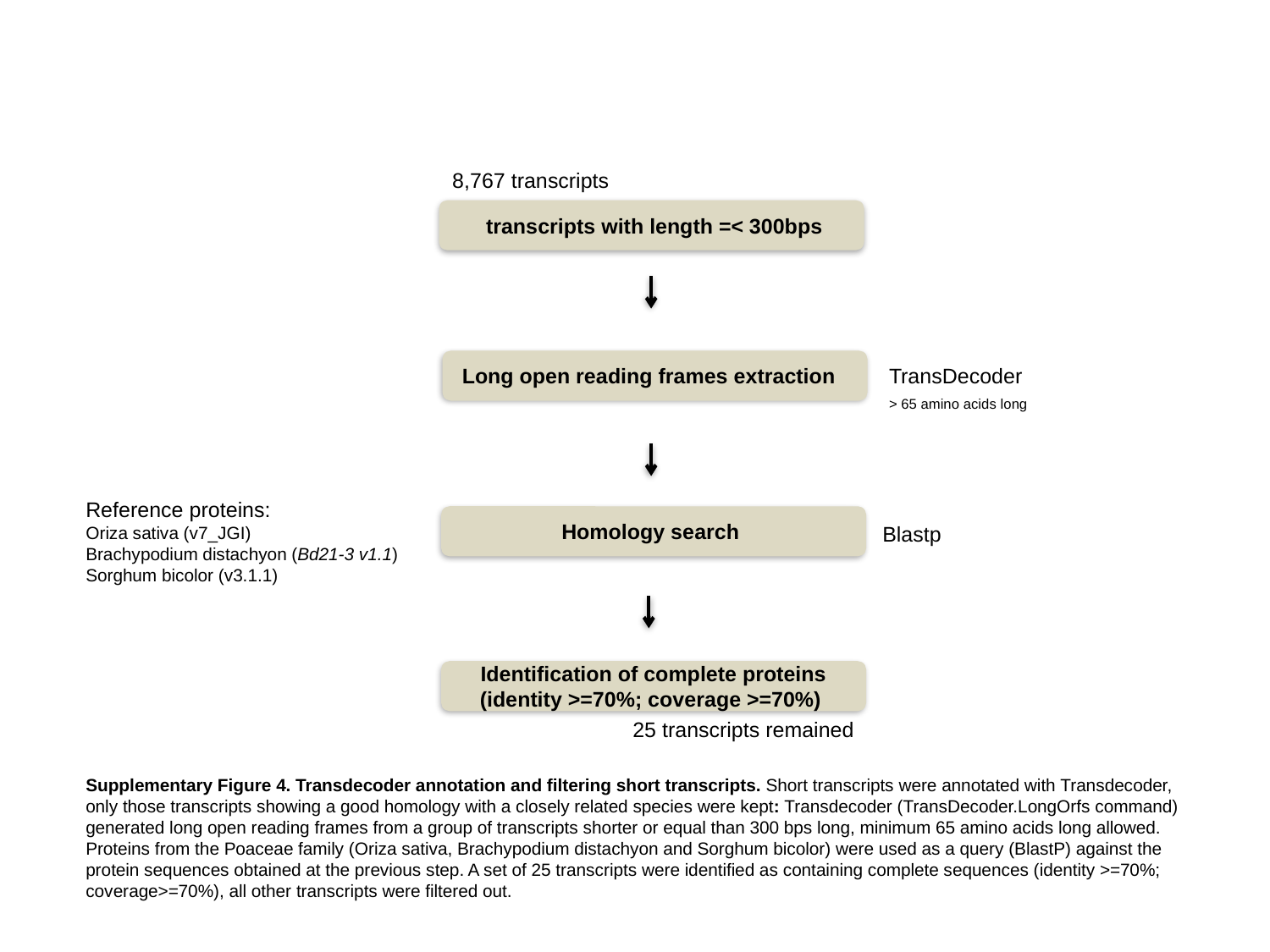

8,767 transcripts
 transcripts with length =< 300bps
Long open reading frames extraction
TransDecoder
> 65 amino acids long
Reference proteins:
Oriza sativa (v7_JGI)
Brachypodium distachyon (Bd21-3 v1.1)
Sorghum bicolor (v3.1.1)
Homology search
Blastp
Identification of complete proteins
(identity >=70%; coverage >=70%)
25 transcripts remained
Supplementary Figure 4. Transdecoder annotation and filtering short transcripts. Short transcripts were annotated with Transdecoder, only those transcripts showing a good homology with a closely related species were kept: Transdecoder (TransDecoder.LongOrfs command) generated long open reading frames from a group of transcripts shorter or equal than 300 bps long, minimum 65 amino acids long allowed. Proteins from the Poaceae family (Oriza sativa, Brachypodium distachyon and Sorghum bicolor) were used as a query (BlastP) against the protein sequences obtained at the previous step. A set of 25 transcripts were identified as containing complete sequences (identity >=70%; coverage>=70%), all other transcripts were filtered out.

### Slide 5
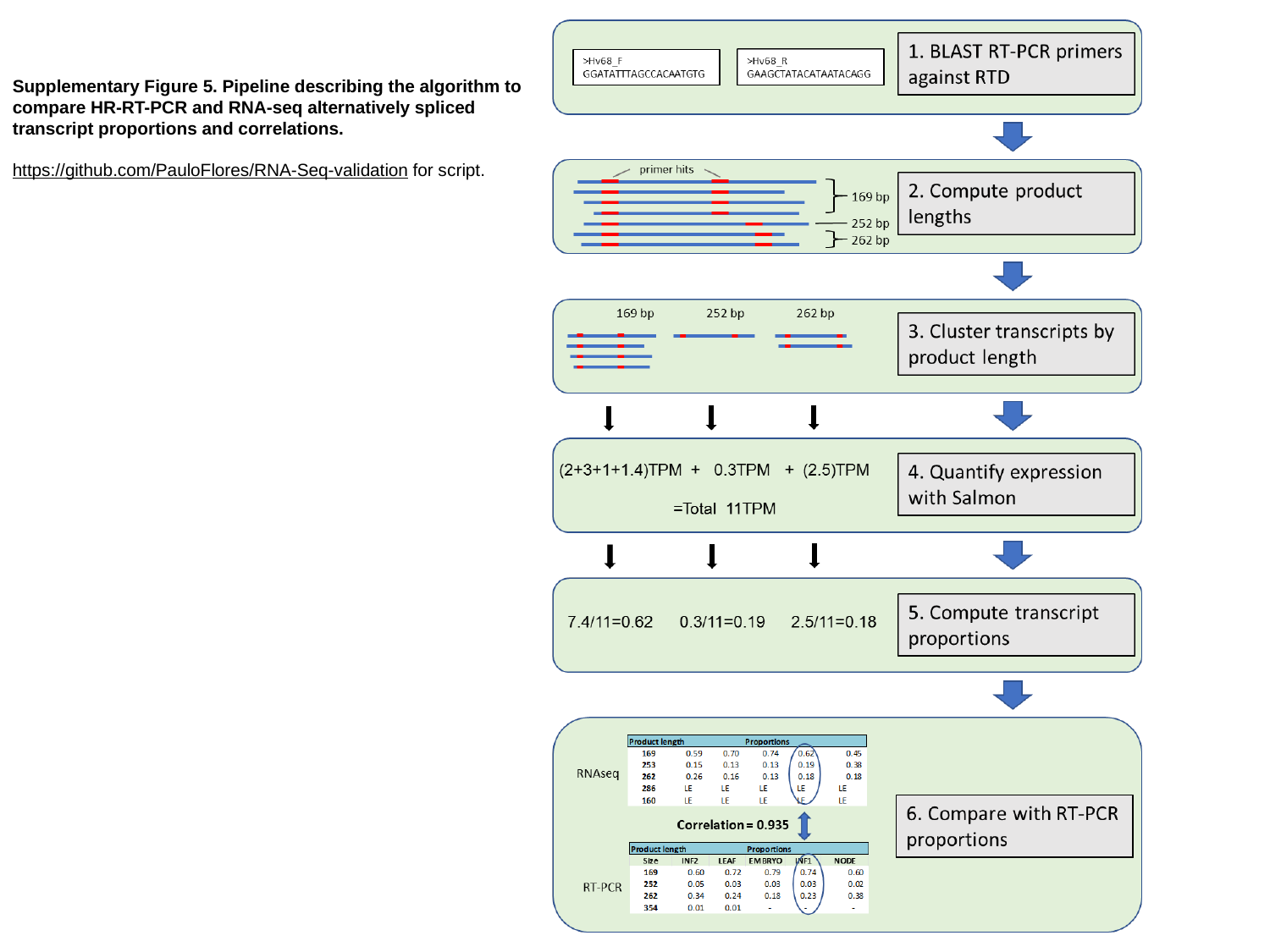

Supplementary Figure 5. Pipeline describing the algorithm to compare HR-RT-PCR and RNA-seq alternatively spliced transcript proportions and correlations.
https://github.com/PauloFlores/RNA-Seq-validation for script.
